# Bayesian Fluorescence Framework for integrative modeling of biomolecules

**DOI:** 10.1101/2023.10.26.564048

**Authors:** Thomas-Otavio Peulen, Andrej Sali

## Abstract

Fluorescence spectroscopic and imaging techniques, such as fluorescence-correlation spectroscopy, image correlation spectroscopy, time-resolved fluorescence spectroscopy, and intensity-based spectroscopy, can provide sparse time-dependent positional and inter-fluorophore distance information for macromolecules and their complexes *in vitro* and in living cells. Here, we formulated a Bayesian framework for processing and using the fluorescence data for interpreting by static and dynamic models of biomolecules. We introduce *Bayesian Fluorescence Framework* (BFF) as part of the open-source *Integrative Modeling Platform* (IMP) software environment, facilitating the development of modeling protocols based in part on fluorescence data. BFF improves the accuracy, precision, and completeness of the resulting models by formulating the modeling problem as a sampling problem dependent on general and flexible libraries of (*i*) atomic and coarse-grained molecular representations of single-state models, multi-state models, and dynamic processes, (*ii*) Bayesian data likelihoods and priors, as well as (*iii*) sampling schemes. To illustrate the framework, we apply it to a sample synthetic single-molecule FRET dataset of the human transglutaminase 2. We show how to integrate time-resolved fluorescence intensities, fluorescence correlation spectroscopy curves, and fluorescence anisotropies to simultaneously resolve dynamic structures, state populations, and molecular kinetics. As BFF is part of IMP, fluorescence data can be easily integrated with other data types to solve challenging modeling problems.

**Statement of Significance:** *Bayesian Framework for Fluorescence* (BFF) is software that implements a probabilistic framework for processing experimental fluorescence data to provide input information for Bayesian integrative structure modeling. BFF facilitates constructing integrative modeling protocols based in part on fluorescence data by reducing the required fluorescence spectroscopy and microscopy domain knowledge. In addition, it improves the precision and accuracy of the resulting models.

## Introduction

In its infancy in the 1950’s, structural biology relied on relatively low-resolution X-ray data and theoretical considerations to unravel the double helical structure of DNA^1^ and the near-atomic structure of myoglobin^2^. Subsequently, thousands of other systems were characterized by X-ray crystallography at atomic resolution^3^. In the 1970’s, X-ray crystallography was joined by nuclear magnetic resonance (NMR) spectroscopy and, in the 2010’s, by 3D electron microscopy (3DEM). Biomolecular structural models of increasing size and complexity have been essential for our understanding of the cellular processes involving the modeled systems.^4^ However, no single experimental technique can currently map all processes involving proteins and their complexes at all size and time scales necessary to understand the studied process.^4,5^ Therefore, to overcome the limitations of individual experimental techniques, it is desirable to compute models by integrative modeling that combines different types of information about the system of interest, including both varied experimental data and prior models.^5,6,7,8^

Modeling is a process that produces models by satisfying input information. Like any modeling, integrative modeling iterates through four stages^4,8^: (*i*) gathering and processing input information; (*ii*) translating information into a representation of the model and a scoring function for ranking alternative models; (*iii*) searching for good-scoring models; and (*iv*) validating models. The input information includes both varied experimental data and prior models (physical theories, statistical analyses, and other prior models). A model representation is defined by the degrees of freedom whose values are computed by modeling (*eg*, Cartesian coordinates of each atom in the case of X-ray crystallography). The scoring quantifies the match of a model to input information; the best scoring function is a Bayesian posterior model density, generally consisting of a product of data likelihoods and priors. The search for good-scoring models may rely on enumeration, optimization, or sampling; Bayesian scoring functions need to be sampled, in principle producing a representative ensemble of sufficiently good-scoring models. Validation provides an estimate of model uncertainty and is essential for a proper interpretation of the model; in Bayesian modeling, the sampled posterior model density is the best such uncertainty estimate. The accuracy of a model is the difference between the model and the truth; the precision (uncertainty) of a model is defined by the spread of the Bayesian posterior model density. The accuracy and precision of the model are determined by the accuracy, precision, and amount of the input information as well as the modeling method that converts input information into a model. The latter, in turn, depends on the model representation, scoring function, and model search. Both model representation and scoring function impact on how precisely a model can reproduce the input information. The computational efficiency of the model search scheme (*eg,* sampling of the Bayesian posterior model density in the case of Bayesian approaches) impacts on its ability to find good-scoring models.

Here, we introduce *Bayesian Fluorescence Framework* (BFF) and the corresponding software that facilitate all steps of integrative modeling based on fluorescence information. BFF acts as a probabilistic preprocessor and abstraction layer that enables integrative modeling by reducing the required domain knowledge in the field of fluorescence spectroscopy and microscopy. We motivate BFF by briefly reviewing fluorescence as an information source that can help reveal the spatiotemporal organization of molecular assemblies in living cells across broad size and time scales. Fluorescence can be a valuable source of information for integrative modeling for six reasons. First, Förster resonance energy transfer (FRET) and photoinduced electron transfer (PET) measurements serve as a spectroscopic molecular ruler to map a distance between fluorophores in the range from 1 to 10 nm with an Ångstrom resolution^9–13^. Second, using fluorescence correlation spectroscopy (FCS)^14^ can map distances as a function of time over 10 decades in time with sub-nanosecond resolution^15^. Third, fluorescence experiments are sufficiently sensitive^16–18^ to study single biomolecules^19,20^. Fourth, fluorescence microscopy can be used to study large biomolecular assembly structures, either *in vitro* or in living cells.^21–26^ Fifth, fluorescence can be measured for systems of nearly any size, ranging from polyproline oligomers^9,27–29^ to ribosomes^30^. Finally, fluorescence measurements can be done in high-throughput mode (*eg*, molecular interactions can be mapped in many live cells).^31^ As a result of the above advantages, fluorescence spectroscopy and microscopy are “bread-and-butter” techniques for studying static and dynamic systems in molecular biophysics. In contrast, they are not established as a standard tool in structural biology. At least in part, this relative lack of use is due to the complexity of the data and the degree of processing needed to accurately recover the distance and kinetic information encoded in the observable dimensions of fluorescence.

Fluorescence experiments, used to characterize FRET, can provide at least eight observables: excitation spectra, fluorescence spectra, anisotropies, fluorescence lifetimes, fluorescence quantum yields, as well as macroscopic time and the fluorescence intensities influenced by the stoichiometry and distance between fluorophores^32–34^. These observables are often processed into distance information^9–13^, which, in turn, could be used to restrain integrative models. Fluorescence spectroscopy observations have already been used to inform molecular models. For example, pioneering theoretical considerations of the fluorophore dipolar coupling (FRET) were combined with sparse experimental data on anisotropy and fluorescence intensities to study chlorophyll^35^. Subsequently, fluorescence information was combined with other data (*eg*, SAXS, EPR, and NMR^36–39)^ to compute models of linear, bulged, kinked, and branched DNA^10,40–45^, RNA and ribozymes^46–48^, protein-ligand complexes^49^, protein nucleic acid complexes^50–57^, dynamic proteins^36,58–62^, large molecular assemblies^30,63,64^, unfolded states of proteins^65^, and intrinsically disordered proteins^66–71^. The studied systems were often represented as rigid bodies^50,57,60,66,67,72,73^ at atomic or coarse-grained resolution^57,74^ in modeling software for X-ray crystallography^72,75–78^, comparative protein structure modeling^79,80^, *ab initio* structure prediction^81–83^, molecular docking^38,84^, and integrative structure modeling^85^. Software optimized for FRET-based modeling, which allows for explicit representation of fluorophores, was also developed.^54,57,74,86^. The explicit fluorophore representation facilitates scoring a model by fluorescence observables. Computationally efficient dye models are used, although they are currently limited to rigid body representations.^54,57^ The position of dyes were modeled by atomistic molecular dynamics (MD)^42,72^, atomistic and coarse-grained Monte Carlo (MC) simulations^24,87,88^, and geometric search algorithms^57^. Dye models were used to map the positional^89^ and orientational distribution of mobile fluorescent proteins^24,64,87,90^ as well as positions of fixed fluorophores in nucleic acids^91^, semi-flexible tethered dyes^54,86^, and flexible tethered dyes.^57,74,92^ Collectively, these studies showed that the dye models are essential for accurate scoring of models informed by fluorescence data.

Summing up, fluorescence-based integrative modeling requires: (a) either domain knowledge for processing raw data or accurate derived distance information, (b) efficient dye models for fast structure searches, (c) an accurate treatment of data and model uncertainties. To fulfill these requirements in a general-purpose integrative modeling platform, we designed and implemented BFF as part of our open-source integrative modeling software IMP environment.^85^ A key design feature of IMP is that it allows mixing-and-matching of different model representations, scoring functions, and model sampling schemes, followed by executing various protocols for model validation and analysis. We also demonstrated its use by application to more than 40 biological systems^7,64,93–143^. IMP allows representing molecules at multiple resolutions, using spatial restraints from many types of data, and searching for solutions by a variety of sampling algorithms. So far, it has been applied mostly to data from EM,^144–153^ small angle X-ray scattering (SAXS),^154–157^ Förster Resonance Energy Transfer (FRET),^158,159^ cysteine and chemical cross-linking,^109,122–124,130,132^ HDX,^135,160^ as well as various proteomics methods.^7,115,161–163^ Next, we describe the framework and its implementation (Methods), illustrate it by ^135,160^a using synthetic data for TG2, the human transglutaminase 2, as practical example (Results), and discuss its implications (Discussion).

## Methods

### BFF

We use Bayes theorem to solve the integrative modeling problem. Bayes theorem quantifies the posterior probability density of model *M*, given experimental evidence *D* and prior information *I*:

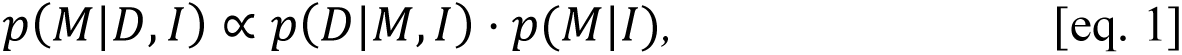

where *p*(*D*|*M*, *I*) is the data likelihood and *p*(*M*|*I*) is the prior. In biomolecular integrative modeling, *M* can include structural, *S*, and dynamic, *K*, parameters, as well as nuisance parameters (*eg*, for backgrounds of signals). *D* is usually experimental information (here, fluorescence data), and *I* is often the prior information, potentially including a molecular mechanics force field and other prior models, such as statistical potentials. The prior expresses the knowledge about *M* before considering *D* (*eg*, assumptions on the background contributions to fluorescence intensities). The data likelihood measures the agreement of the model with the data. Computing the data likelihood requires a forward model and a noise model. The forward model predicts *D* for *M*, assuming no experimental noise. The noise model quantifies the distribution of the differences between *D* and the forward model^164^, which originates from the random experimental error, systematic experimental error, and error in the forward model; noise models generally assume that systematic and forward model errors are 0. In time-resolved fluorescence spectroscopy, the forward model computes model fluorescence decay curves. The noise model considers the Poissonian counting statistics of the photon data and is combined with the forward model into a data likelihood function.^165^

Integrative modeling problems can be solved in a single step using a complex likelihood function that combines all information. In fluorescence, information on a structure is composed of multiple datasets of different kinds, each having its own nuisance parameters. Thus, the dimensionality of *M* to be computed by integrative modeling grows when considering additional data. This increase in dimensionality is an issue, as searching for models consistent with the data usually requires a larger computational effort for higher dimensional models. For integrative models of T4 lysozyme, we previously described time-resolved FRET experiments for 33 FRET pairs.^166^ A measurement of a FRET pair consists of two fluorescence decay curves. Each curve requires at least 4 nuisance parameters for the fluorescence background, the instrument response function (IRF) background, the IRF time-shift, and the scattered light fraction. Thus, a model for all measurements contains at least 264 nuisance parameters. We previously sampled these model parameters by a semi-automated global analysis approach.^166^ In this approach, the dimensionality is reduced by introducing relations among parameters from different datasets.^167^ In addition to estimating the parameters (*eg*, inter-dye distance), we also estimated their uncertainties arising from data noise and other sources, using traditional error propagation ^166^. Despite the success of this approach, three challenges remain: achieve scaling with the size of the experimental data, introduce a formal framework for combining varied data, and estimating uncertainty of nonlinear parameters.

In BFF, we address these challenges by a divide-and-conquer approach, breaking down the higher dimensional problem into multiple lower dimensional problems that can be sampled more easily, followed by coupling the optimal and suboptimal solutions of the lower dimensional problems to obtain a complete solution. We formulate this divide-and-conquer approach using probabilistic graphical models (PGMs)^168^ to encode a Bayesian hierarchical posterior model density. A PGM is a graph that encodes conditional dependencies among random variables described by probability density functions (pdfs). Here, the random variables correspond to the BFF parameters. Bayesian hierarchical models are layered statistical models that estimate model parameters of posterior distributions using the Bayesian method. The output values of the BFF parameters are then translated into spatial restraints for Bayesian integrative structural modeling (**Fig 1**).

**Figure 1.**
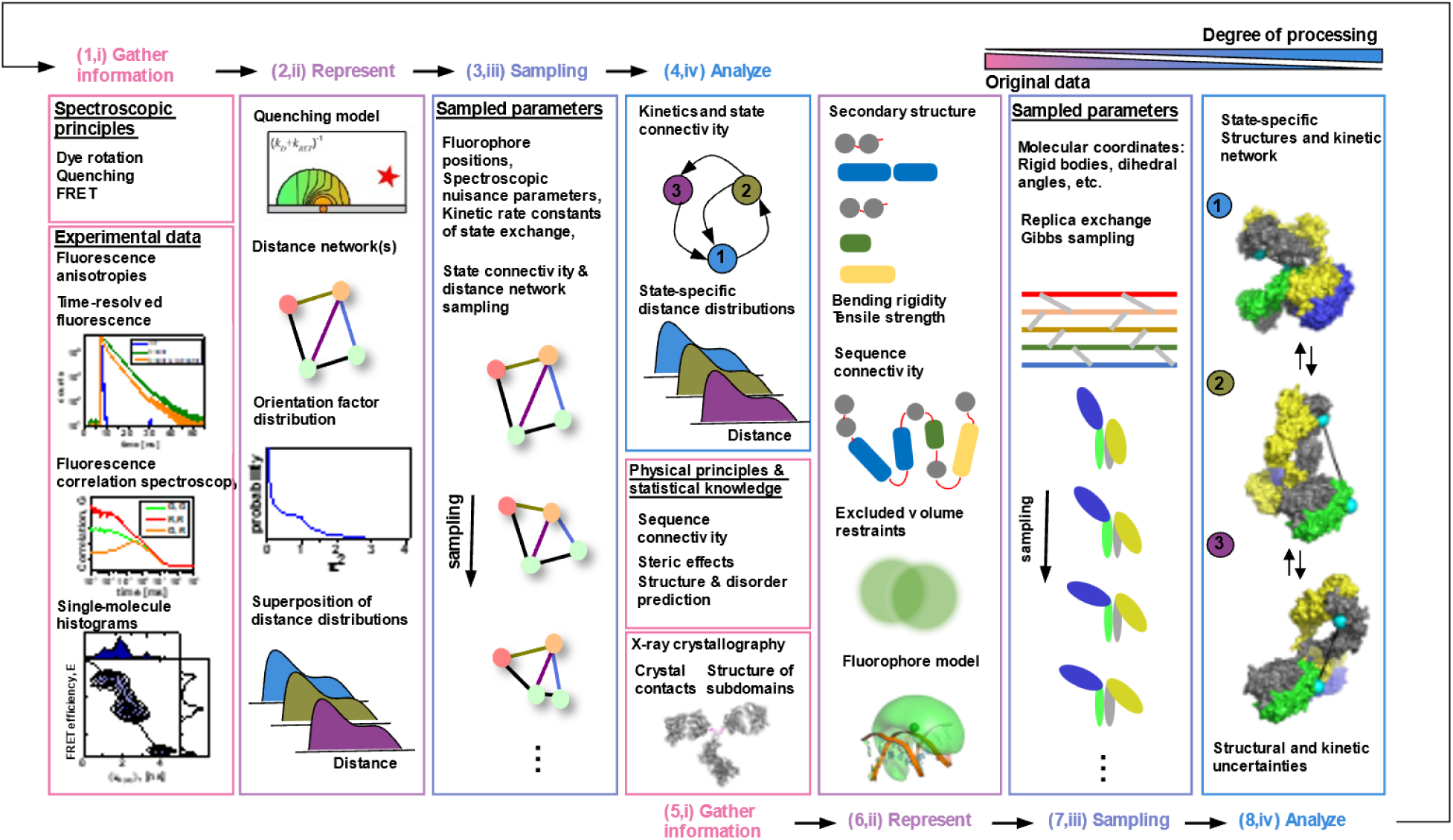
The Bayesian fluorescence framework (BFF) feeds information into the integrative modeling platforms to combine multiple input sources for dynamic structure biology. BFF uses a Bayesian approach that considers multiple data sources with associated data noises to maximize the information that is robustly transformed into an output. We feed BFF information into the integrative modeling platform (IMP). We model systems using BFF and IMP in four stages where we: (*i*) gather all information about the system that we wish to include and decide at which stage of modeling the information is applied; (*ii*) define representations for the input information including the degrees of freedom we wish to assess; (*iii*) sample the representation parameters; (*iv*) process and analyze the sampled parameters. BFF converts experimental fluorescence data into spatial and kinetic information that abstracts experimental nuisances for outputs that can be used in restraints for molecular integrative modeling. At stage (1*, i*) we gather multiple experimental data such as fluorescence intensities, time-resolved fluorescence identities, fluorescence correlation spectroscopy curves, single-molecule counting histograms, fluorescence anisotropies, and information on fluorescence quenching. At stage (2, *ii*) we define representation for the data that produces that serve as spatial or kinetic input for later stages. At stage (3, *ii*) the parameters of the representations are sampled and processed at stage (4, *iv*) to yield an input for later stages. At stage (5, *i*) we combine the output of stage (4, *iv*) with additional molecular information such as structural information provided by other data sources. At stage (6, *ii*) we define molecular representations that we sampled in stage (7, *iii*) and analyzed in stage (8, *iv*) using IMP.

### Modeling structure and structural kinetics of transglutaminase 2

We illustrate BFF by a synthetic benchmark, corresponding to modeling the known structures and structural kinetics of human transglutaminase 2 (TG2) based on synthetic fluorescence data. TG2 consists of three domains whose configuration changes upon binding of ATP.^169^ To model TG2, we define a hierarchical posterior model density with two independent layers. In the first layer, BFF combines likelihoods and priors for fluorescence data. The output of the first layer is a model posterior density for the distance and kinetic information (**Fig.1**, top). The second layer solves the structural modeling problem using BFF outputs and additional structural information as an input for standard integrative structure modeling with IMP (**Fig.1**, bottom). The fluorescence modeling and structural modeling layer are each an integrative modeling problem whose solution can be described in terms of four stages^4,8^ (**Fig.1**). Next, we describe these 8 stages.

#### Data acquisition, stage 1

In the first BFF stage, we begin by gathering fluorescence data (**Fig.1**). For TG2, this data consists of simulated photon traces from single-pair FRET (spFRET) experiments. In a spFRET experiment, a single donor, D, and acceptor fluorophore, A, are attached to the molecule of interest at defined residue positions. By construction, two TG2 conformers, C_1_ and C_2_, are in fast exchange (**Fig.2**). As structures for C_1_ and C_2_, we chose the monomeric subunits of crystallographic dimers 3ly6^170^ and 2q3z^171^, respectively. A dynamic exchange between C_1_ and C_2_ was modeled with typical but arbitrarily defined values of the first order rate constant in each direction. The fluorophores in a FRET pair were attached to TG2 by flexible linkers with a length of 20 Å and had a Förster radius, *R*_0_, of 52 Å. To pick the D and A attachment sites, we use an automated planning approach that selects an optimal set of FRET pairs by maximizing the information from FRET experiments^74^; an optimal set of FRET pairs maximizes the precision of a structural model ensemble that satisfies the corresponding distance restraints (and other given information). We selected a set of FRET pairs for modeling structures and an additional set of FRET pairs for validating structures. For each FRET pair ensemble, we simulated time-resolved fluorescence decays and single-molecule multiparameter fluorescence detection experiments.^32,33^ To mimic typical experiments, we use experimental anisotropy data^166^ as templates for the simulations. The FRET-pair photon traces were processed to produce fluorescence decays, *F*, time-resolved anisotropies, *r*, and FCS curves, *G* (**Fig.3A**). *F*, r, and *G* contain information on distances and population, dipole orientations, and molecular kinetics, respectively, and serve as the input fluorescence data for BFF.

**Figure 2.**
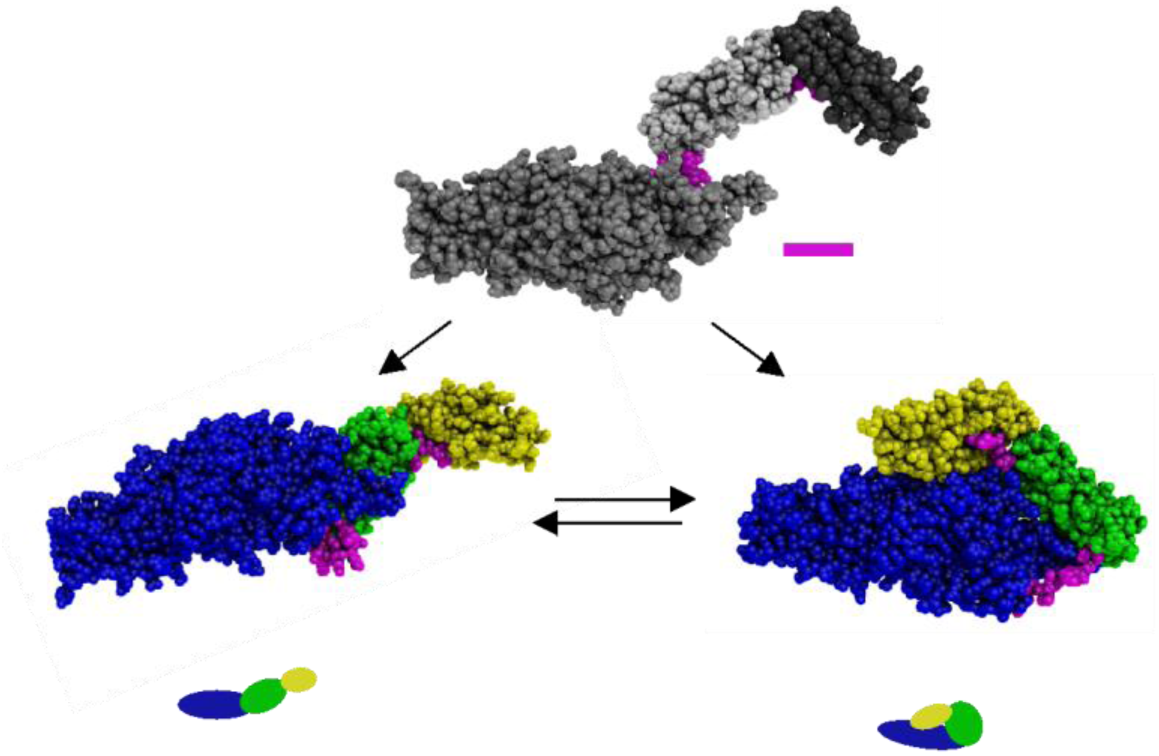
Presentation of the test case. We use the human transglutaminase 2 (TG2) to introduce the Bayesian Fluorescence Framework (BFF). For crystal structures of TG2 of the open (bottom left) and the closed (bottom right) conformation we generate synthetic single-molecule time-resolved FRET experiments. We referred to these conformations as *C_1_* and *C_2_*, respectively. In the simulations *C_1_* and *C_2_* are in a dynamic exchange. To simultaneously resolve the structures of *C_1_* and *C_2_* and the rate constants by BFF, we combine BFF with the integrative modeling platform (IMP). In IMP we represent TG2 by three domains that are linked by flexible regions (magenta spheres). The amino acids sequence numbers (aa) of the corresponding domains are presented next to the protein representation of *C_2_*. Starting from randomized initial configurations (top) we find posterior distributions of the structures and exchange rate constants for *C_1_* and *C_2_*.

**Figure 3.**
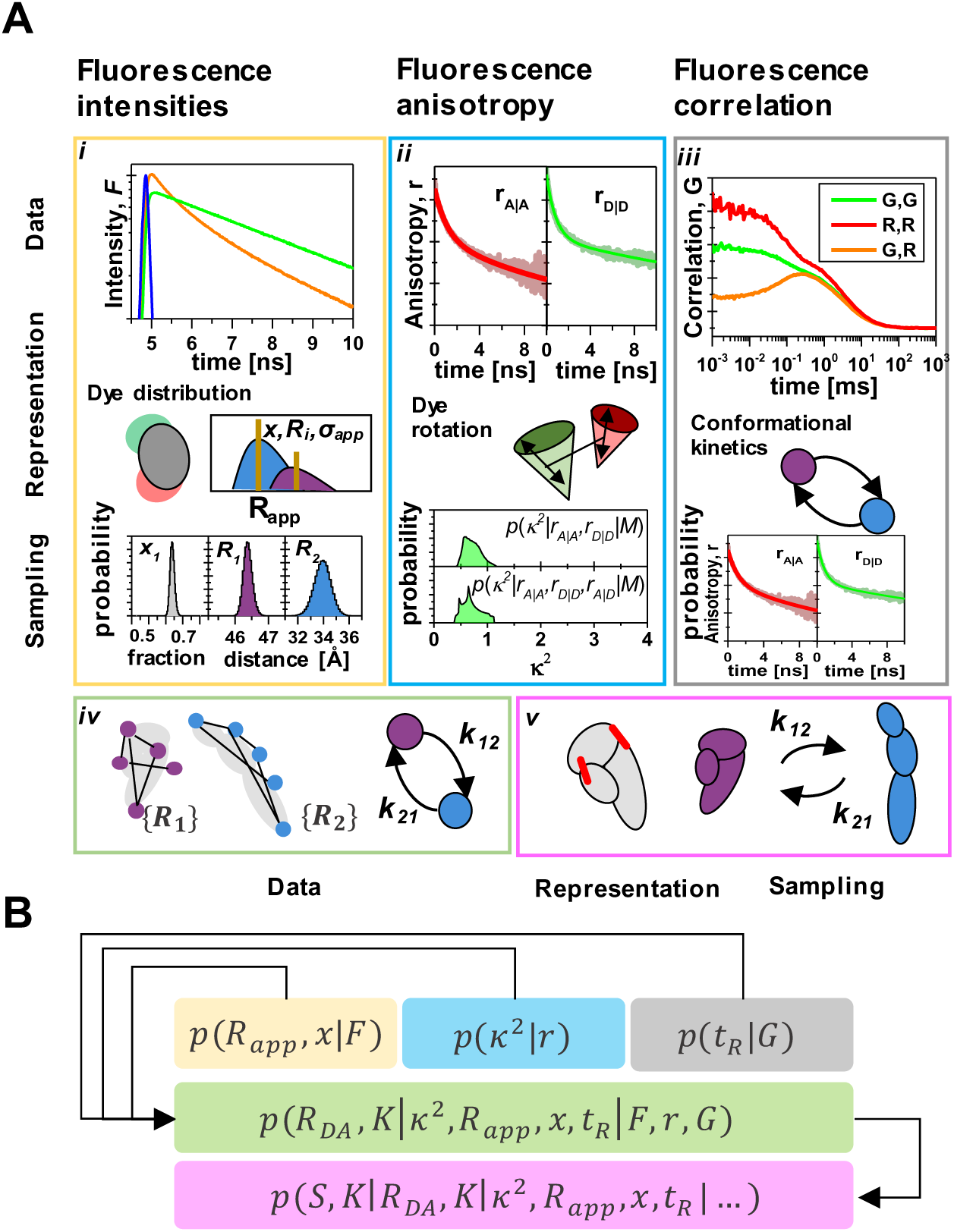
“Divide-and-conquer” approach of the Bayesian fluorescence framework (BFF). (**A**) BFF processes time-resolved fluorescence intensities (yellow), fluorescence anisotropy (blue), and fluorescence correlation data (gray). The posterior combining the fluorescence data recovers distance networks for states and associated exchange rate constants (green) that serve as input for the integrative modeling platform (IMP) to yield dynamic structures (magenta). BFF combines data, data noise, representation of the data, and sampling of representation parameters to yield posterior distributions. (*i*) Time-resolved fluorescence intensities decays (yellow box) in the presence of FRET inform on apparent inter-dye distance distributions. The data is represented considering prior knowledge on the spatial distribution of the dyes around their attachment site (representation) to determine posteriors of inter-dye distances of states and populations (sampling). (*ii*) Fluorescence anisotropy decays (blue box) inform on the dye rotation and the relative orientation of the donor and acceptor dipoles. Here, the dipole orientations follow a wobbling in a cone (WIC) model to yield probability distributions of the orientation factor, *k*^2^, consistent with the input information, i.e., the residual anisotropies, *r*_∞_, of the directly excited donor (D|D) and the acceptor (A|A) (top green distribution) and of the FRET sensitized acceptor that is excited via FRET (A|D) (bottom distribution). (*iii*) Fluorescence correlation spectroscopy (FCS) curves (gray box) inform on dynamics. Here, FCS curves were represented by a kinetic model with a discrete number of states and observables parameters (diffusion time, correlation amplitudes, relaxation times) were sampled. The posterior distributions of the relaxation times, *t*_*R*_, characterize the kinetic network. (*iv*) The information is gathered (green box) to yield distance networks and exchange rate constants assembled in a rate matrix, *K*, and (*v*) used for Bayesian structural modeling in IMP (magenta box) with molecular representations. (**B**) Illustration of the probabilistic model that combines fluorescence intensities, *F*, FCS, *G*, fluorescence anisotropy information, *r*, to recover structures, *S*, and transition rate matrix, *K*. Fluorescence intensities, *F*, (yellow) yield posteriors of apparent distances, *R*_*app*_, and population fractions, *x*. Fluorescence anisotropies, *r*, inform on the posterior of the orientation factor, *k*^2^ (blue) and FCS curves on posteriors of relaxation times, *t*_*R*_. The combined posterior distributions yield distances for states of a corresponding transition rate matrix, *K* (green). The assembled information recovers structures, *S*, that are in dynamic exchange as described by *K*.

#### IMP-BFF, stage 2

In the second IMP-BFF stage, we define representations and data likelihoods to convert the input fluorescence data *F*, *r*, and *G* into spatial and kinetic features of the modeled system, including apparent distances, orientation factor distributions, and relaxation time distributions, respectively. *F* informs distributions of apparent distances between labeled sites, *p*(*R*_*app*_|*F*) (**Fig.3A**,*i*). In a FRET experiment with a donor, D, and an acceptor, A, *F* depends on the inter-dye distance, *R_DA_*, and the relative orientation of the fluorophore transition dipole moment, expressed by the orientation factor *κ*^2^ (**Fig.3A**,*ii*).^172^ Thus, *F* only informs on apparent distances, *R*_*app*_, if *k*^2^ is not considered. *r* informs on the rotational mobility of fluorophores and can be used for *κ*^2^ inferences, *p*(*κ*^2^|*r*).^173^ *G* informs on molecular relaxation times^15,174^, *p*(*t*_*R*_ |*G*) (**Fig.3A**,*iii*).

To convert *F* into apparent distances, we describe fluorescence decays in the absence and the presence of FRET by a joint forward model that combines multiple independent time-resolved fluorescence experiments.^175^ The forward model computes fluorescence decays and depends on nuisance parameters and on a distribution over species fraction, *x*(*R*_*app*_).^175^ We determine the posterior *p*(*x*(*R*_*app*_)|*F*, *M*_*R*_) for individual FRET pairs by sampling to find *x*(*R*_*app*_) that reflects both the input *F* and the prior *p*(*x*(*R*_*app*_), *M*_*R*_) (**Fig.3A**,*i*, top). We parametrize and sample from a *x*(*R*_*app*_) using a model *M*_*R*_ with a defined number of protein conformational states and consider the conformational heterogeneity associated with the fluorophore linker. *x*(*R*_*app*_) models are distributions composed of sub-distributions. The sub-distributions reflect protein conformational states. As previously described,^172,176^ the width and amplitude of the sub-distributions relate to the conformational heterogeneity of the fluorophore and the population of the protein states, respectively (**Fig.3A**,*i*, middle). We score the fluorescence decays considering the photon shot-noise^165^.

To convert *r* into orientation factor distributions, we represent the orientation factor distributions, *p(k^2^|M_k_*, *r*), by a orientation factor model *M*_*κ*_(**Fig.3A**, *ii*). As *M*_*κ*_, we use a wobbling-in-a-cone (WIC) model ^177^. We use the residual anisotropies of the directly excited donor, D, *r*_*D*|*D*_,∞, and the directly excited, *r_A|A,∞_*, and FRET sensitized, *r_A|D_*,∞, acceptor, A, as anisotropy information, *r* (**Fig.3A**, *ii*, top). We score *p*(*k*^2^|*M*_*k*_, *r*) considering uncertainties, Δ*r*_∞_.

To convert *G* into relaxation time distributions, we process *G* for each FRET pair individually. The *G* of each FRET pair consists of a set of time-dependent auto (ACFs) and cross-correlation functions (CCFs) determined by correlating the signal intensities by color FCS (cFCS) (**Fig.3A**, *iii*, top). We represent the correlation functions by a product of a diffusion and a kinetic term for the molecular diffusion and kinetics, respectively. The kinetic term contains relaxation times and corresponding correlation amplitudes as parameters (**Supplementary Note1**). The relaxation times are the inverse eigenvalues of a matrix specifying the reaction rate constants, thus describing the kinetics of the process. The correlation amplitudes are affected by various factors, such as background signals, spectral cross talks, and molecular brightness ^15,174^. We describe *G* of each FRET pair by a forward model that includes the kinetic terms of the ACF and CCFs (**Fig.3A**, *iii*, middle). We score models by χ^2^ (*ie*, a data noise-weighted sum of squared deviations of the model and the data). The data noise is approximated by the standard deviation of multiple experimental correlation curves.

#### IMP-BFF, stage 3

In the third BFF stage, we convert *F*, *r*, and *G* into posterior distributions of spatial and kinetic features of the modeled system by sampling over the unknown parameters. We determine posteriors of *p*(*x*(*R*_*app*_)|*F*) for each FRET pair individually (**Fig.3A**, *i*, bottom) using a Markov Chain Monte Carlo (MCMC) approach.^178^ To determine *p*(*k*^2^|*r*, *M*_*k*_), we sample over the dipole orientations consistent with the assumed (*i*) *M*_*k*_ model as well as (*ii*) the experimental residual anisotropies and their uncertainties for each FRET pair (**Fig.3A**, *ii*, bottom).^172^ We sample over all FCS parameters, including the kinetic FCS terms. We also obtain probability distributions of relaxation times, *p*(*t*_*R*_|*G*) (**Fig.3A**, *iii*, bottom) using the MCMC approach.^178^

#### IMP-BFF, stage 4

In the fourth BFF stage, we analyze and couple model posterior densities. Inspired by recently described Bayesian metamodeling, we call a model posterior density of model *X* conditioned on data Y, *p*(*X*|*Y*), a surrogate model of *Y*. We call the process of combining surrogate models coupling. We couple surrogate models by additional conditional probability distributions, called couplers. Here, we use physics-based couplers that express theories and physical laws in terms of probability distributions of coupled variables. We couple surrogate models to produce spatial and temporal information that can be interpreted without considering the original fluorescence data. This way we create input restraints for integrative modeling (**Fig.2A**). Specifically, we couple here surrogate models of *F*, *r*, and *G* (**Fig.3A**, *i*-*iii*) to obtain physical inter-dye distances, *R*_*DA*_, and molecular rate constants (**Fig.3A**, *iv*), as follows.

First, we couple surrogate models of *F* and *r* to infer inter-dye distances, *R*_*DA*_, to resolve the issue that *F* depends on *R*_*DA*_ as well as the fluorophore rotation^175,179,180^ and translation ^177,181,182^. We combine information on *R*_*app*_ and *k*^2^:

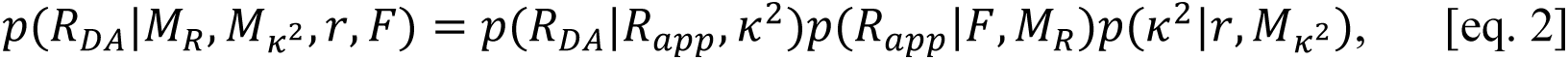

where *p*(*R*_*DA*_|*R*_*app*_, *κ*^2^) is a coupler that relates *R*_*app*_, *k*^2^, and *R*_*DA*_. We simulated fluorophores that rotate fast compared to the FRET process. For such cases, following Förster’s theory, *R*_*DA*_ = (3 ⁄ 2 ⋅ *k*^2^)^1⁄6^*R*_*app*_ . Thus, *p*(*R*_*DA*_|*R*_*app*_, *κ*^2^) = *δ*(*R*_*DA*_ − (3 ⁄ 2 ⋅ *k*^2^)^1⁄6^*R*_*app*_), where *δ* is the delta function. By conditioning *p*(*R*_*app*_) and *p*(*k*^2^) on *F* and *r*, respectively, we incorporate information about *r* and *F*. In the absence of anisotropy information, *p*(*R*_*app*_) can be converted into *p*(*R*_*DA*_) using an uninformative prior *p*(*κ*^2^). We compute *p*(*R*_*DA*_) for typical *p*(*k*^2^) to illustrate the relation between *p*(*R*_*DA*_) and *p*(*R*_*app*_) (**Supplementary Fig. 2**). Couplers are problem specific. A different coupler is required for fluorophores rotating rapidly^179^ and slowly^24,87,183^ on the time-scale of the FRET process.

Next, we couple *F* and *G* surrogate models to recover first order rate constants. *F* and *G* inform populations and relaxation times, respectively. To combine this information, we define a physical model that relates *p*(*x*(*R*_*app*_)|*F*) and *p*(*t*_*R*_|*G*). To relate *x*(*R*_*app*_) and *t*_*R*_, we describe the evolution of the system. We represent the system by a set of coupled first-order linear differential equations:

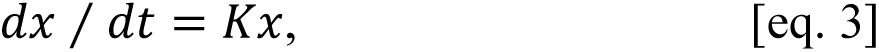

where *K* is a matrix filled with the first order rate constants, and *x* is a vector of the state population. The rate constants describe the time-evolution and thus the steady-state population, *x*, of the states and relaxation time, *t*_*R*_, informed by the fluorescence intensity, *F*, and the fluorescence correlation data, *G*, respectively. We use Bayes theorem to obtain the posterior of *K*:

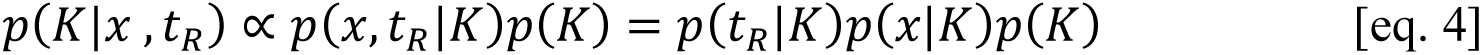

where the factorization is made possible the statistical independence of the parameters *x* and *t*_*R*_. In summary, the description of steps 1-5 above illustrates how BFF facilitates the factorization of the posterior model density into likelihoods and priors, based on a physical understanding of the relationships between the data, model, and modeling variables.

#### IMP-Structure, stages 5 to 8

Structural modeling requires a molecular representation, a scoring function, and a structural sampling scheme. The goal of sampling is to find structural models that are consistent with the input information as quantified by the scoring function. To achieve this goal here, we use our open-source *Integrative Modeling Platform* (IMP) package (https://integrativemodeling.org) ^4,8,85^. The molecular representation specifies Cartesian coordinates of the protein, including labels. The scoring function includes terms for excluded volume and FRET (**Supplement Note 2**). The sampling scheme includes a rapidly exploring random tree algorithm (RRT)^184^ and MCMC simulations (**Supplementary Fig. 5**). All the input data files scripts and output files are freely available (**Data and Software Availability**).

## Results

### Modeling structure and structural kinetics of transglutaminase 2

We now describe modeling of structure and structural kinetics of transglutaminase 2, discussing the results from each of the 8 stages of the entire process (Methods): stage 1 (Data acquisition), stages 2-4 (IMP-BFF), and stages 5-8 (IMP-Structure).

#### Data acquisition

Similarly to the planning stage of a typical FRET project, we selected FRET pairs and simulated the corresponding synthetic experiments (stage 1). In the planning stage, we assumed an uncertainty of 6% in the FRET efficiency and selected a set of 11 informative FRET pairs (155-680, 388-588, 366-502, 498-661, 406-523, 465-592, 425-540, 550-659, 425-519, 363-661, and 406-610) and a set of 6 additional pairs (173-659, 155-571, 406-661, 425-646, 385-661, and 406-618) for modeling and validation, respectively. For the 11 FRET pairs, we expect to find a structure with an average *C*_α_-RMSD of 2.5 Å (**Supplementary Fig. 1**).

#### IMP-BFF

We processed the data from simulated experiments by IMP-BFF to compute distances for structural modeling by IMP. To obtain these apparent distances, we modeled *F* by a two-state representation, with apparent distances *R*_*app*,1_and *R*_*app*,2_and associated population fractions *x*_1_ = *x* and *x*_2_ = 1 − *x* for the states *C_1_* and *C_2_* ^175,176^. We computed *p*(*R*_*app*,1_, *R*_*app*,2_, *x*|*F*) by sampling (**Fig. 3**). To convert apparent distances to inter-dye distances for structural modeling, we modeled the orientation factor *k*^2^ using anisotropy information, *r* (**Fig. 3**). We coupled probability densities of *k*^2^and apparent distances to obtain inter-dye distances. The ground truth for the distances, probability densities for apparent distances, and inter-dye distances are visualized in **Supplementary Fig. 3**. The average deviation between the *R*_*DA*_ground truth for C_1_ and C_2_ is –2.9 Å and 0.25 Å, respectively. The average *R*_*DA*_ root-mean-squared-deviations (RMSD) between the reference and mean for C_1_ and C_2_ are 6.0 Å and 5.9 Å, respectively. The average *R*_*DA*_ precision for C_1_ and C_2_ are 8.8 Å and 9.4 Å, respectively. The *R*_*DA*_precision is lower than the precision assumed in the FRET pair search. Thus, we expect to recover structures at a resolution worse than 2.5 Å RMSD.

To determine posteriors of unimolecular rate constants that describe the exchange between conformations states, we couple posterior distributions of species population fractions, *p*(*x*|*F*), to posterior distributions of relaxation times, *p*(*t*_*R*_|*G*). For an individual dataset *F*_*i*_, we found C_1_ species population fraction, *x*, in the range of 54% to 78%, with uncertainties in the range of 0.5% to 9.9%. The combined posterior of the species fraction for all FRET-pairs 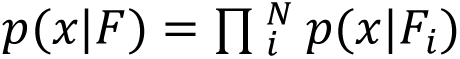 is 66.6 ± 0.3%. This combined posterior is closer to the reference value of 66.6% than any individual *p*(*F*_*i*_) (**Fig. S4**) and has a significantly higher precision. We used a representation that jointly describes the auto and cross-correlation functions of color FCS experiments and outputs relaxation times (**Supplementary Note 1**). A a two-state system is described by a single relaxation time, *t*_*R*_. We sampled the FCS model parameters consistent with the input data and determined the posterior model density of the relaxation time (**Fig. 3A**,*iii*). To yield the probabilities of the reaction rate constants, we coupled *p*(*G*) with *p*(*x*|*F*). For the test case, the rate constants *k*_12_ for *C*_1_ → *C*_2_and *k*_21_ for *C*_2_ → *C*_1_ relate to the C_1_ species fraction, *x*_1_ = *k*_21_ ⁄ (*k*_12_ + *k*_21_), and the relaxation time *t*_*R*_ = *k*_12_ ⁄ (*k*_12_ + *k*_21_). The combined probability of *p*(*k*_12_, *k*_21_|*x*) and *p*(*k*_12_, *k*_21_|*t*_*R*_), *p*(*k*_12_, *k*_21_|*x*, *t*_*R*_), agrees with the ground truth (**Fig. 4**).

**Figure 4.**
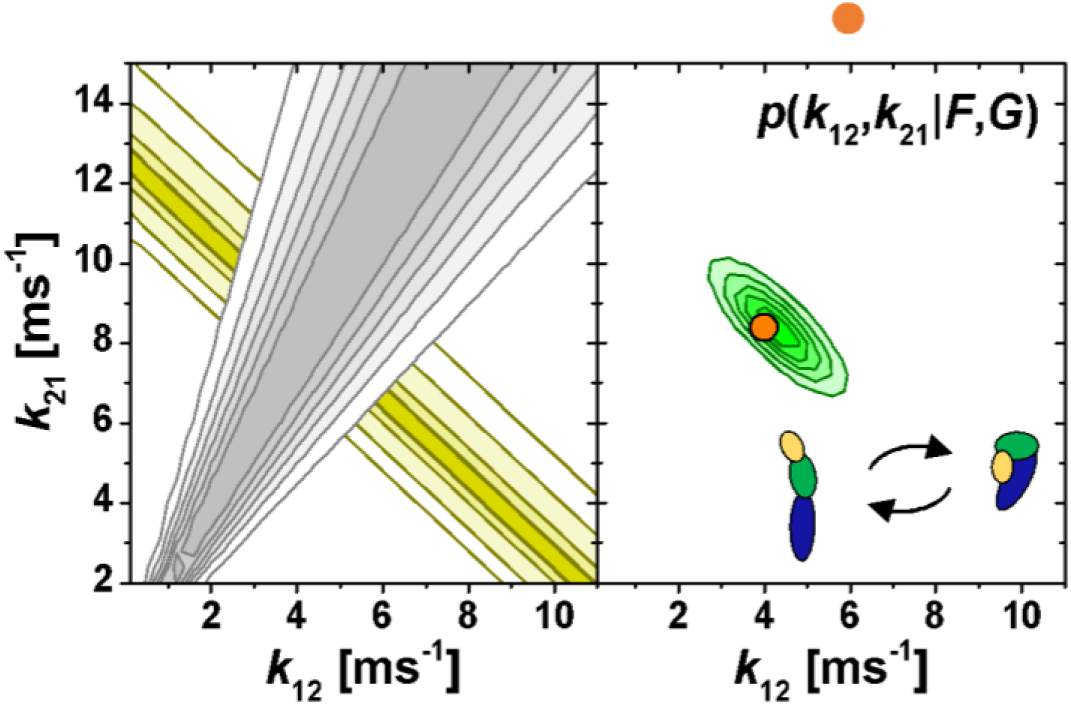
Kinetic posterior distributions recovered by the Bayesian Fluorescence Framework (BFF) for synthetic single-molecule FRET (smFRET) experiments of the human transglutaminase 2 (TG2) test case. The posterior distribution for the rate constants *k*_12_ and *k*_21_ that describe the exchange between the conformers C1 and C2. To the left the posterior distribution for the fluorescence intensities *p*(*k*_12_, *k*_21_|*F*) (yellow) and the correlation functions *p*(*k*_12_, *k*_21_|*G*) (gray) are shown. To the right the combined posterior distribution *p*(*k*_12_, *k*_21_|*F*, *G*) (green) is displayed. The circle marks the pair of rate constants (*k*_12_, *k*_21_) that was used to generate the synthetic smFRET experiments.

#### IMP-Structure

We now use the probability densities of inter-dye distances to find structures for conformations C_1_ and C_2_ (**Fig. 2**). The conformations used to generate the synthetic data serve as a ground truth reference to assess the model accuracy. The initial structure together with the distance information obtained for the fluorescence data (**Fig. 1**, step 5i) was used to generate structures using a forward model for flexible dyes and a scoring function implemented in IMP (**Supplementary Note 2**), using a workflow outlined in **Supplementary Fig. 5**. The output posterior model densities for C_1_ and C_2_ correspond to structures consistent with the data used for modeling. We quantify the structure model precision by the average Cα RMSD. The posteriors of C_1_ and C_2_ have an average Cα RMSD of 10.0 Å and 6.9 Å, respectively (**Fig. 5**, **Supplementary Fig. 6**). The precision of the recovered C_1_ and C_2_ is considerably lower than the average precision of 2.1 Å anticipated in the planning stage (**Supplementary Fig. 1**). In the planning stage, we sampled the representation independent of the experimental data, selected a set of FRET pairs that optimize a single state model selection problem, and used a simplistic noise model. Therefore, a lower precision was expected. The accuracy of the structures best describing the C_1_ and C_2_ in the sampled posterior as quantified by the deviation from the ground truth is 16.3 Å and 7.6 Å in Cα RMSD from C_1_ and C_2_, respectively (**Fig. 5, Supplementary Fig. 6**).

**Figure 5.**
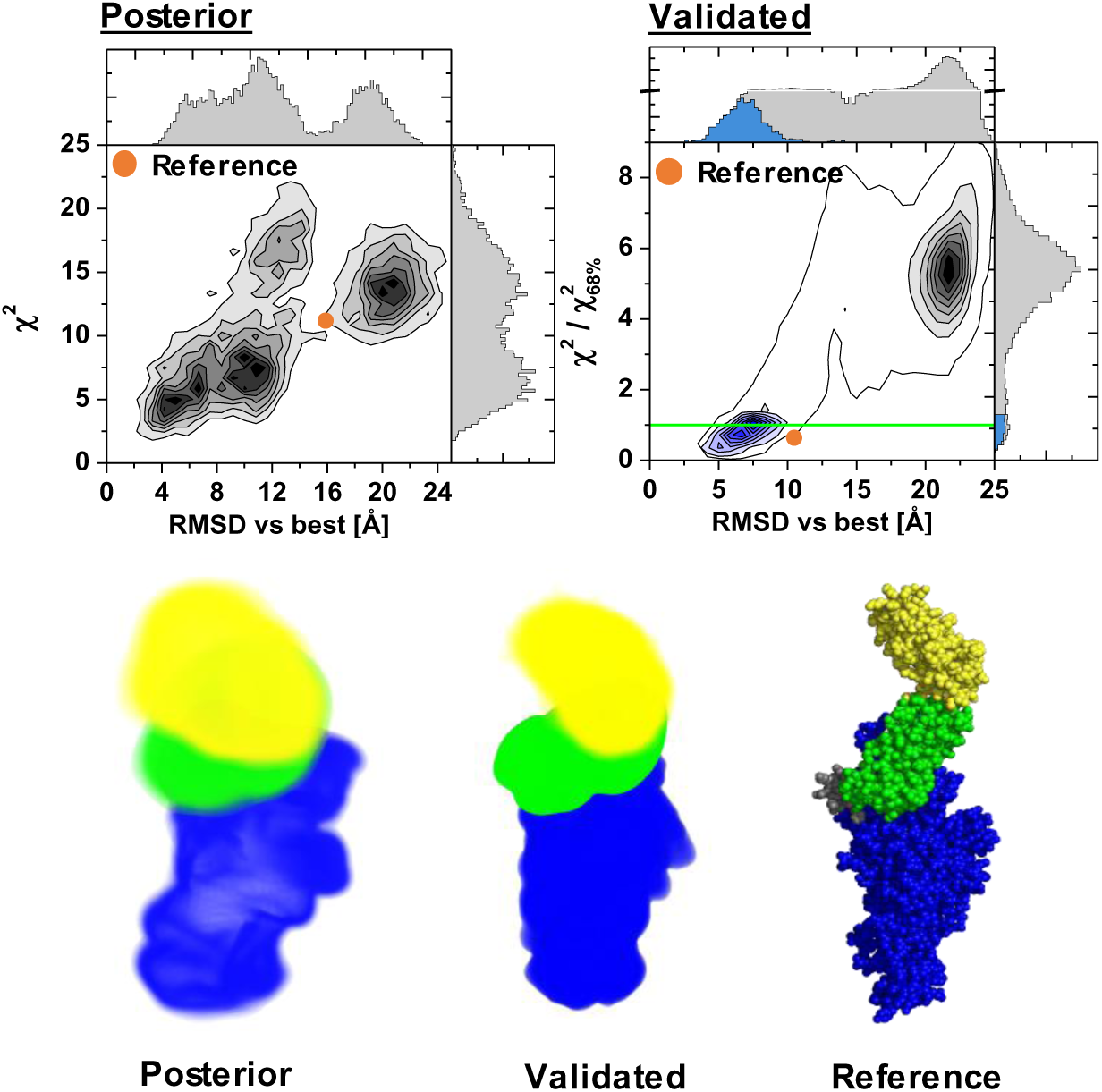
Conformational posterior distributions and validated structures for C_1_ recovered using Bayesian Fluorescence Framework (BFF) outputs and the Integrative Modeling Platform (IMP) for the human transglutaminase 2 (TG2) test-case. The top left panel is a frequency histogram of the C_1_ posterior distribution for the sum of uncertainty weighted squared deviations, χ^2^, vs. the *Cα* RMSD for the best structure. The top right displays for C1 the sum of noise weighted squared deviations between the model and data using data not used in the modeling normalized by a confidence level of 68%, χ^2^/χ^2^ . Structures above the green horizontal line are inconsistent with the data not used for modeling at a confidence level of 68%. To the bottom the densities for the posterior model distribution (left) and the validated structures (middle) are displayed. The bottom right displays an atomistic representation of the C_1_ ground truth. The localization densities of the amino acids (aa) 1-460, aa 473-583, and aa 590-688 are colored in blue, green, and yellow, respectively. The orange circles correspond to the ground truth structures were used to generate the synthetic data.

Finally, we validate the structures generated by IMP using data not used for modeling (**Fig. 5, Supplementary Fig. 6**). We use the sum of the uncertainty weighted squared deviations between the model and the data, χ^2^, normalized to a reference χ^2^value for a certain confidence level ^74^. Structures with sum of squared deviations above 1 are inconsistent with the validation data for a defined confidence level. At a 68% confidence level, 6.8% (33,600 out of 489,320) of the C_1_ and 48% (108,560 out of 226,240) of the C_2_ structures in the posterior are consistent with the validation data (**Fig. 5, Supplementary Fig. 6**). In the experiment planning phase, FRET pairs were selected for low redundancy. Thus, it is expected that a large fraction of the structures is discriminated against in the validation stage. Structures best describing the validation data deviate from the C_1_ and C_2_ ground truth by 11.1 Å and 8.1 Å in *C_α_* RMSD, respectively. The larger deviation for C_1_ is anticipated, because the structure used to obtain C_1_ and C_2_ is based on a perturbed closed TG2 conformation and the fine differences between C_1_ and C_2_ cannot be captured by the coarse representation we chose to describe the system. This distinction between C_1_ and C_2_ highlights the value of model validation when designing representations. Moreover, it stresses the need for a versatile modeling platform, such as IMP. In conclusion, the global domain arrangement and the exchange kinetics were accurately captured (**Fig. 5, Supplementary Fig. 6**).

## Discussion

We introduced the Bayesian Fluorescence Framework (BFF) that converts experimental inputs using Bayes theorem to probabilistic outputs for integrative modeling of structure and kinetics. The framework transforms data with associated uncertainties into posterior model densities that can be used as surrogates for the experimental data. We demonstrated the framework on synthetic single-molecule and ensemble fluorescence data. We processed time-resolved fluorescence intensities, fluorescence correlation spectroscopy curves, and fluorescence anisotropies as inputs and showed how to combine information to yield probability densities of rate constant matrices and distances for Bayesian integrative structural modeling. We also computed the corresponding structures and kinetics.

In the FRET field, the joint analysis of multiple linked physical models has a long tradition. Experimental data by a single analysis kernel is referred to as global analysis. In global analysis, a mapping from a target representation with parameters representing physical invariants of the system to the experimental data is defined (forward model) and the parameters are optimized on the multidimensional data surface using physical representations that utilize the implicit relations between the parameters and the data sets^167^. Global analysis optimizes a complex scoring function for all data and reduces the number of free parameters by introducing dependencies among model variables. For large and diverse datasets, this approach becomes computationally unfeasible because (*i*) sampling a high-dimensional scoring function is difficult and (*ii*) multiple programs for different data types need to be closely integrated. Therefore, experimental data (e.g., fluorescence decays, fluorescence anisotropies, or single-molecule counting histograms) has been often analyzed separately, followed by combining point estimates for the parameters and their uncertainties using traditional uncertainty propagation ^166^, although the global analysis approach is more accurate^167^.

BFF is designed to benefit from both global analysis and multiple individual analyses. This goal is achieved by the use of probability densities for model parameters as opposed to point estimates (i.e., particular realizations of model parameters and their uncertainties). This divide-and-conquer approach allows us to partition high-dimensional likelihood functions and priors into parts that can be sampled independently and combined in a second step using probabilistic couplers. As a result, BFF has six advantages. First, nonlinear dependencies among parameters and/or couplers can be accounted for by a rigorous Bayesian treatment of uncertainties. Second, existing analysis methodologies and data can be easily incorporated (demonstrated here for fluorescence anisotropies, fluorescence intensities, and fluorescence correlation curves). Third, the outputs of probabilistic fluorescence analysis software can be mixed-and-matched without the need for a tight integration in a global analysis kernel. Fourth, computing can be decentralized and parallelized using independent modulus. Fifth, modeling can be easily updated each time new data, input model, surrogate model, and/or a coupler is provided. Finally, BFF modeling can potentially be integrated relatively easily into the Bayesian metamodeling framework.^185^

We showed how BFF abstracts complex aspects of fluorescence data to provide an input for integrative modeling of molecular structures and kinetics. BFF defines data sources and inputs for multiple experiments, processing units (e.g., correlators, burst selectors, etc.), and probabilistic data transformations. As a result, BFF helps with using FRET data for Bayesian integrative modeling of structures and kinetics. In addition, the BFF framework may help overcome the lack of standard operating procedures for measurements and analysis^186^ and data exchange formats for interoperable software in FRET analysis^69,186^. We plan to incorporate additional fluorescence data types, such as pulsed interleaved excitation^187^, multi parameter fluorescence detection single-molecule counting histograms^32^, and other forward models and representations,^188,189^ improve the representation and coupling of surrogate models to simplify the use of probabilistic fluorescence data processing, and introduce representations that abstract fluorescence information without the need to specify the number of states. These technical and scientific advances could greatly facilitate the processing of fluorescence data and enable advanced fluorescence experiments as standard tools for integrative modeling of structures and kinetics.

Beyond structural biology, the BFF framework may also contribute to the modeling of cellular processes. The BFF approach could be applied to data from live-cell high resolution imaging of molecular assemblies^190,191^, spectroscopic imaging of metabolic states^192,193^, diffusion and interactions mapping of proteins through spatiotemporal image correlation spectroscopy^194^, raster image correlation spectroscopy^195^ and pair correlation functions^196^ of high-throughput live-cell data^31^, ultimately resulting in integrative models of biomolecular processes in the cell.

## Data and Software Availability

BFF is implemented as a module of our open source *Integrative Modeling Platform* (IMP) software (https://integrativemodeling.org)^8,54^. The source code of the development version of the software is available at (https://github.com/fluorescence-tools/imp.bff). A compiled version of the software and documentation are available at https://anaconda.org/tpeulen and https://docs.peulen.xyz/bff, respectively.

## Supplementary Figures

**Supplementary Figure 1.**
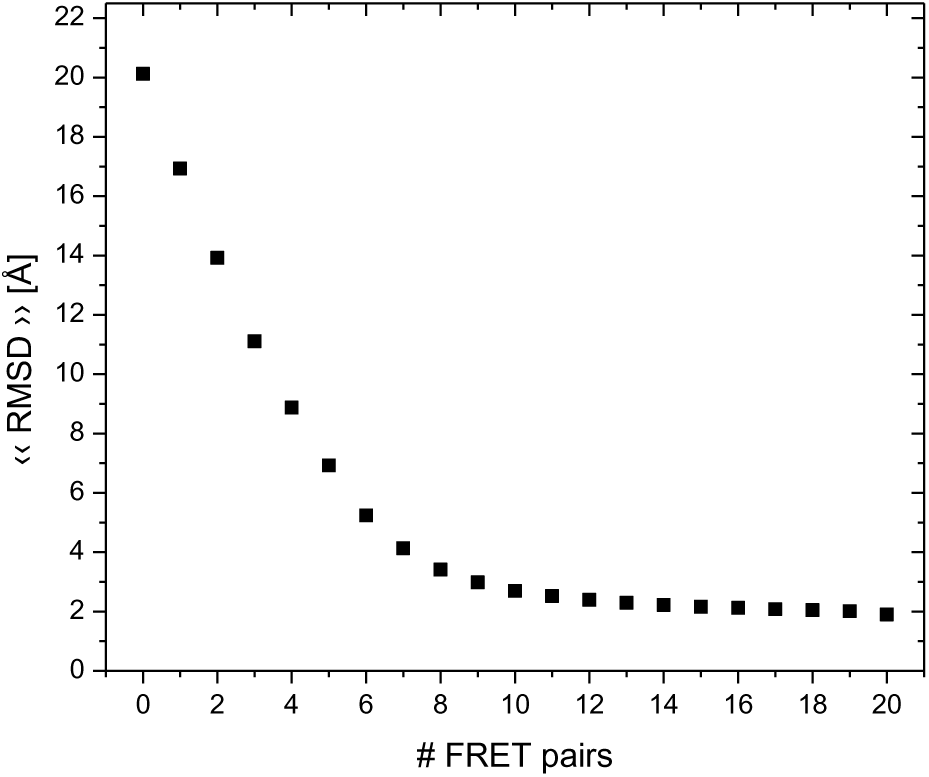
FRET pair selection for the human transglutaminase 2. The average expected root-mean-squared-deviation (RMSD) is plotted in dependence of the number of FRET pairs. The amino acid pairs selected for labeling are in order: 155-680, 388-588, 322-680, 366-502, 498-661, 406-523, 322-502, 465-592, 425-540, 550-659, 425-519, 363-661, 406-610, 173-659, 155-571, 406-661, 425-646, 385-661, 406-618, 222-519.

**Supplementary Figure 2.**
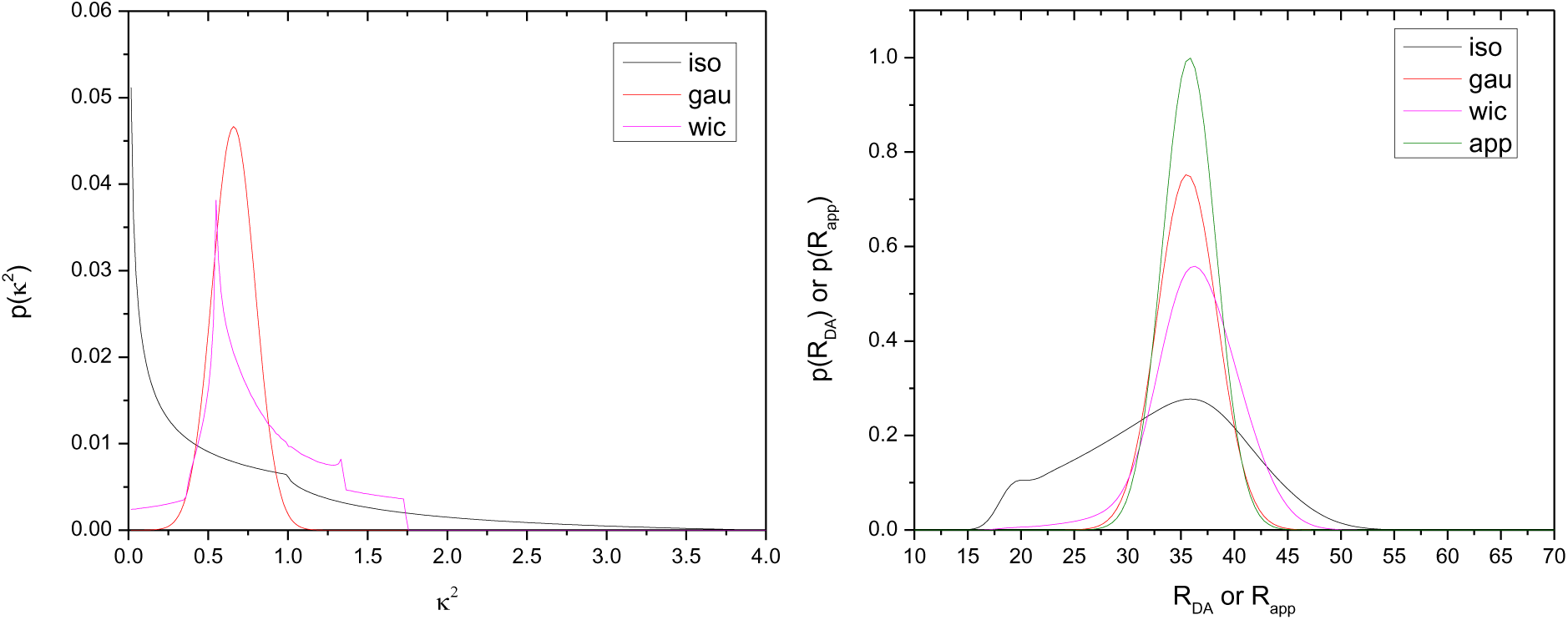
Effect of different orientation factor distributions, *p*(*k*^2^), on inter-dye distance distributions, *p*(*κ*^2^), for *R*_*app*,2_ of the TG2 FRET pair 155-680. Left, input orientation factor distribution *p*(*k*^2^) for an isotropic orientation model (iso), a wobbling-in-a-cone-model (WIC) informed by residual anisotropies, and a normal distributed *p*(*κ*^2^) with an average *κ*^2^ of 0.66 (gau)(). Right, input apparent distance distribution *p*(*R*_*app*_) and output inter-dye distance distributions, *p*(*κ*^2^), for the different *p*(*k*^2^).

**Supplementary Figure 3.**
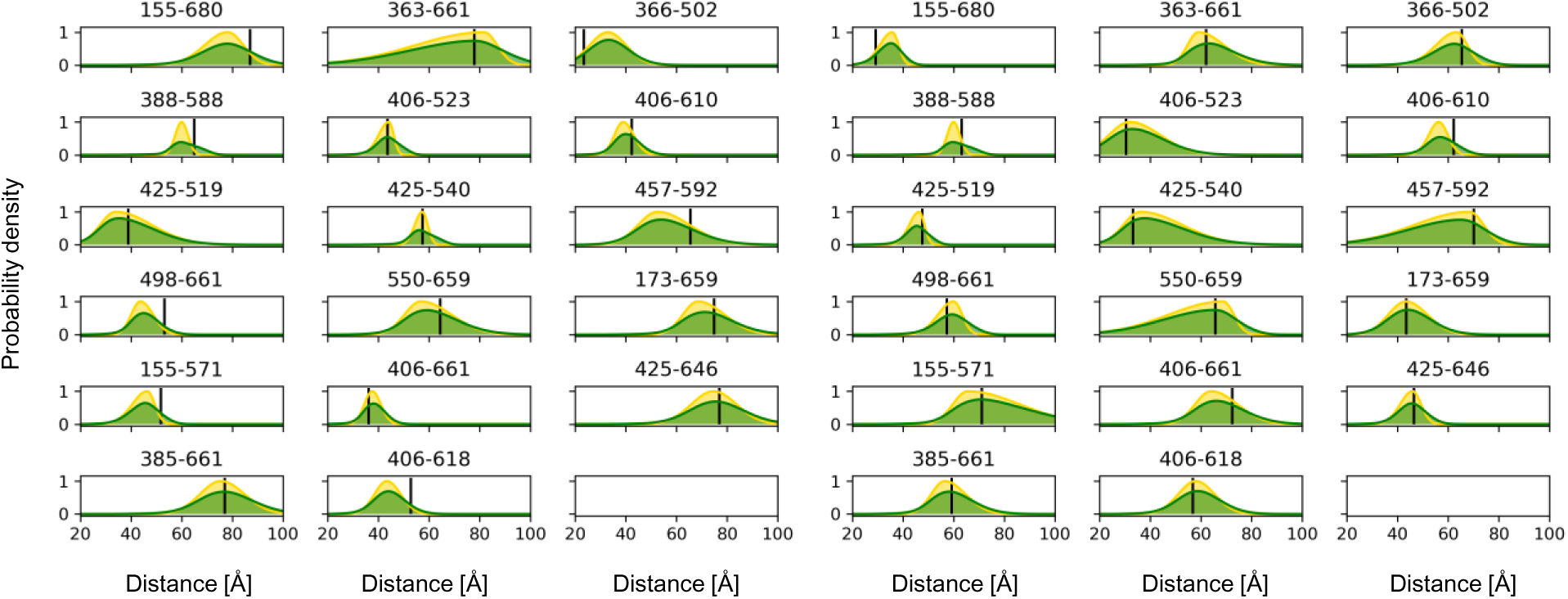
Apparent, inter-dye, and ground truth distances for synthetic TG2 data. The probability distributions of apparent distances, *p*(*R*_*app*_), from a Bayesian analysis of fluorescence decays (yellow). Distributions of the inter-dye distances, *R*_*DA*_, were recovered by combining *p*(*R*_*app*_) with knowledge on the orientation factor, *k*^2^. Average inter-dye distance of all FRET pairs used in modeling and validation for the reference structures with the PDB-ID 2q3z (state C1) and 3ly6 (state C_2_) are shown as black vertical lines. Posterior probability densities of the apparent distances *R*_*app*,1_ and *R*_*app*,2_ (yellow) and the inter-dye distance *R*_*DA*_ (green). The pair of numbers shown to the top of the plots refers to the amino acids in FRET pairs that were used for labeling.

**Supplemental Figure 4.**
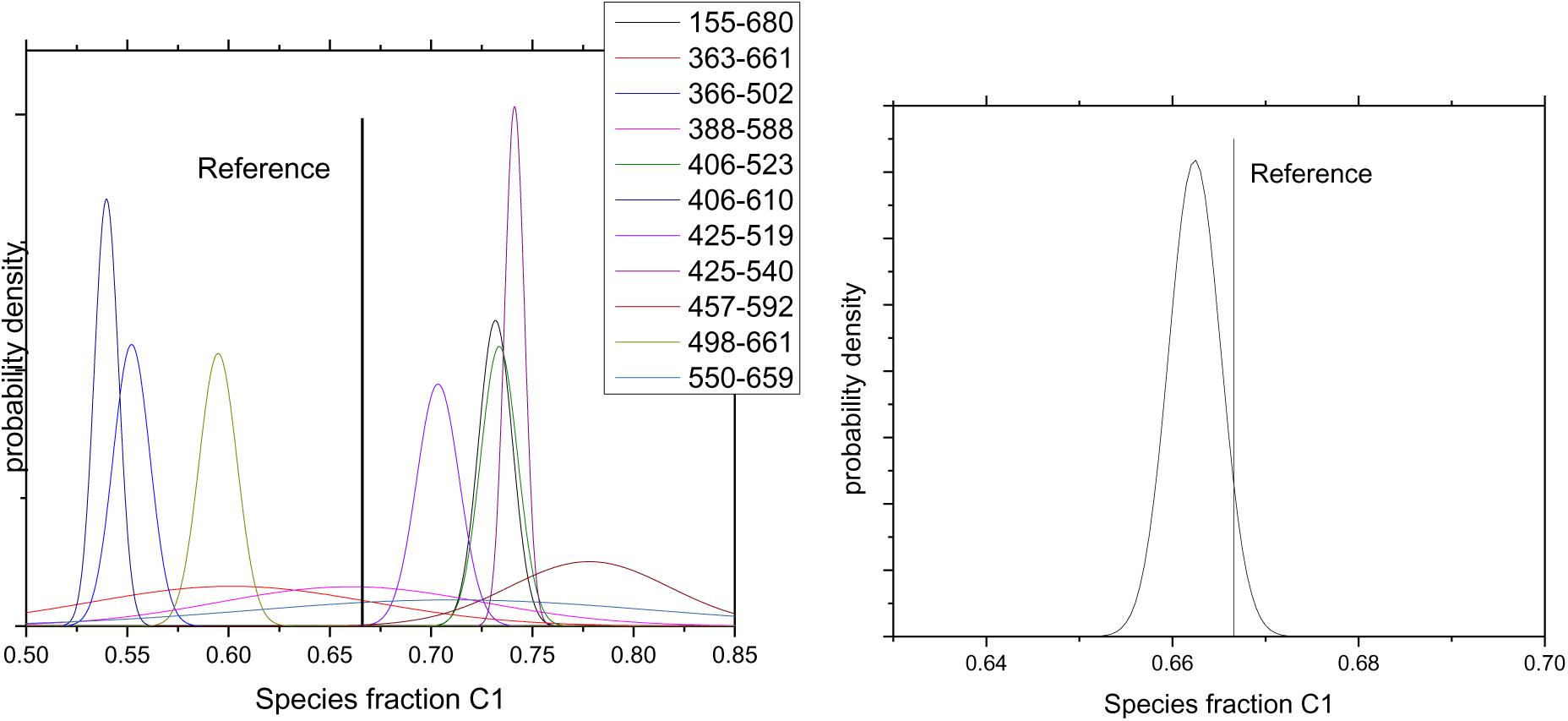
Combining species fractions densities. To left the probability density distributions of the species population fractions of the state C1 for the individual datasets are displayed. To the right the product of all probability densities is displayed. The vertical lines marks the species population fraction of the state C1 that was used to generate the synthetic experiments that serves as ground truth reference.

**Supplementary Figure 5.**
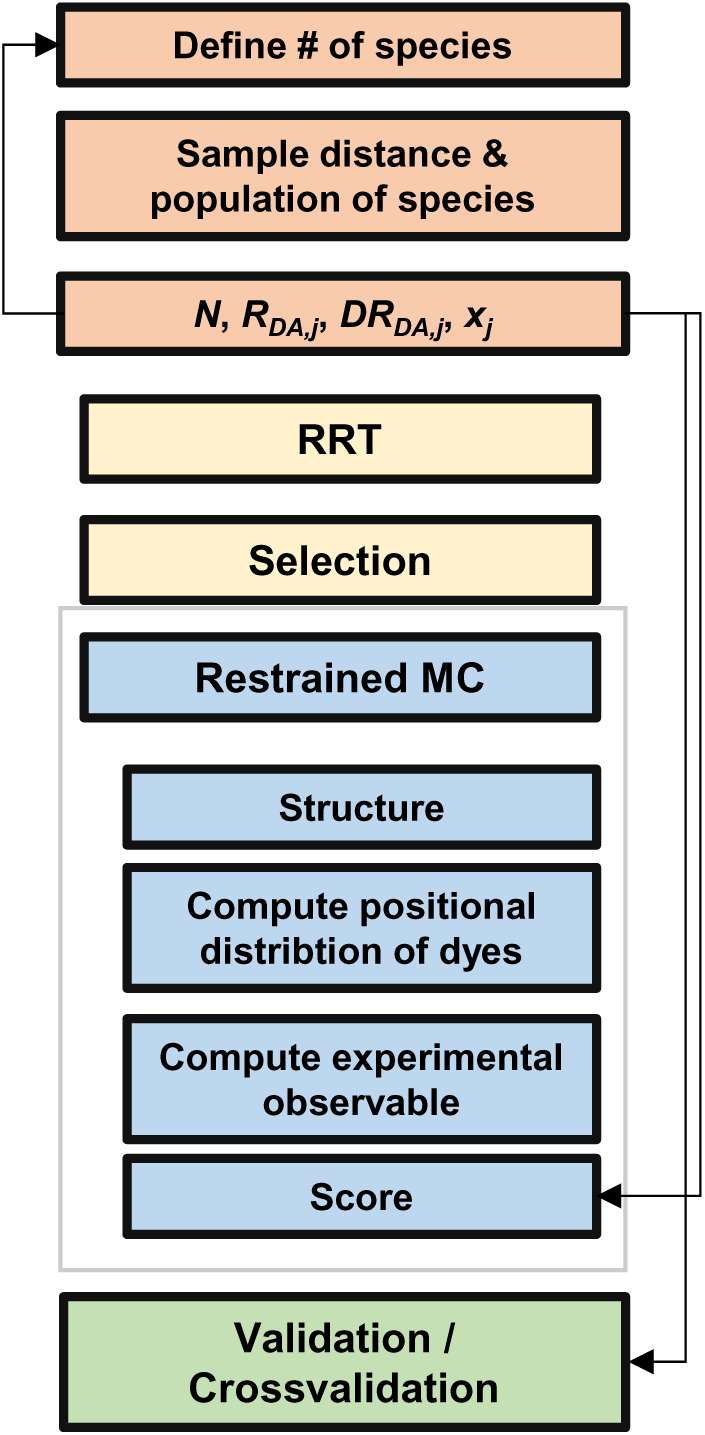
Structural modeling workflow. After defining the number of species (step 1) the probability distribution of species distances and population are recovered (step 2, 3) via BFF. Next, structures are sampled using the representation using the Rapidly exploring Random Tree (RRT) algorithm implemented in the integrative modeling platform (IMP) to generate a set of structures that were used as seeds for FRET-restrained Monte Carlo (MC) sampling. In the FRET-restrained MC for (step 6.1) the positional distribution of the dyes is computed (step 6.2). Next, given the spatial distribution of the dyes experimental observables are computed (step 6.3) to score (step 6.4) the structure from step 6.1. The distance information of the BFF is in part used for scoring and generating new structures (step 6.4) and in part used for validation (step 7).

**Supplementary Figure 6.**
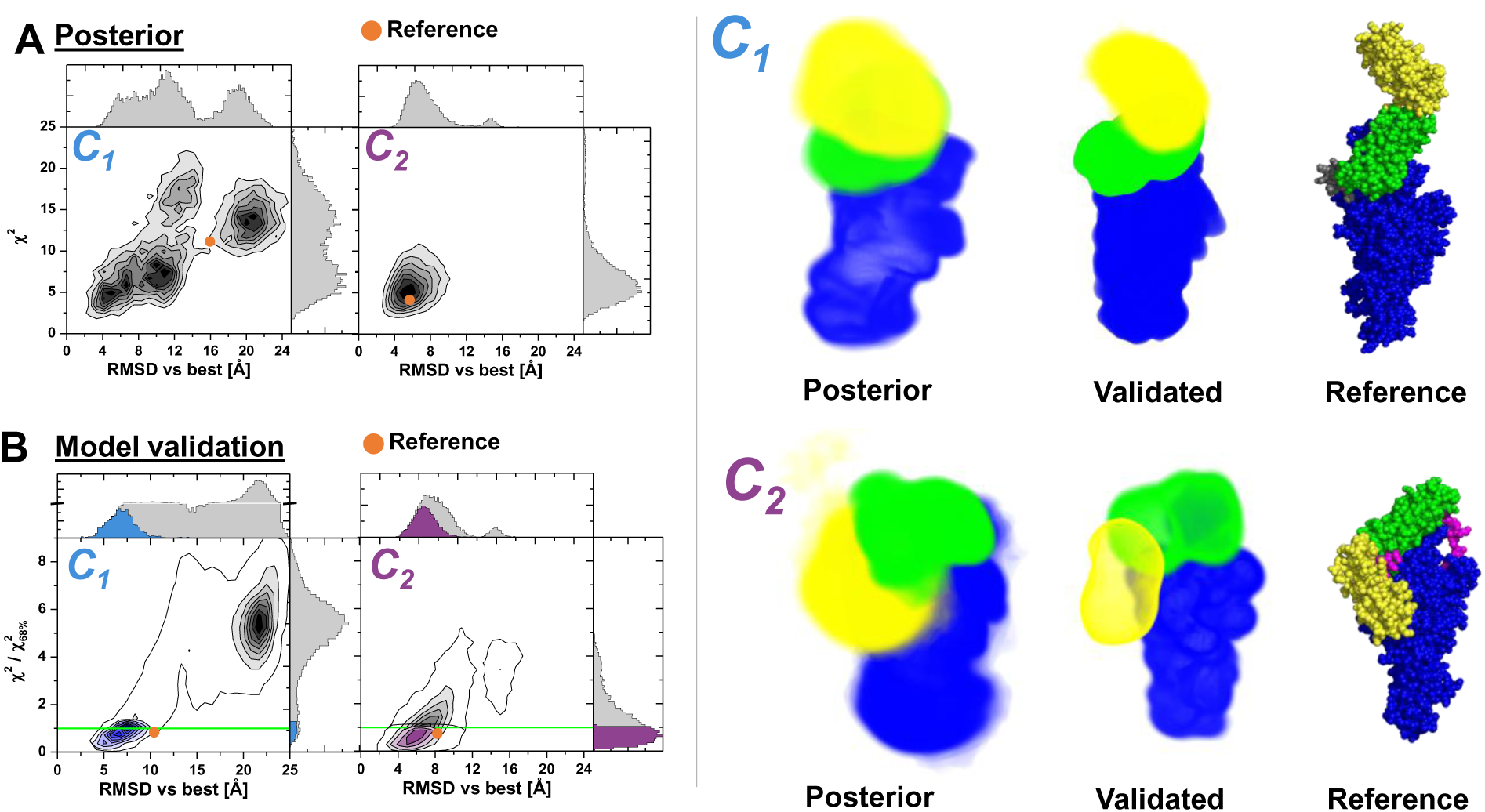
Model posterior densities and validated ensembles. **(a)** Sum of uncertainty weighted squared deviations between model and data, χ^2^, in dependence of the *C*_α_ root-mean-squared-deviation (RMSD) vs the best structure for the conformers C_1_ and C_2_ as recovered by the BFF outputs and integrative modeling via IMP. The orange circle corresponds to the reference structures that were used to generate the synthetic experimental data. (**b**) Histogram over all modeled structures for C_1_ and C_2_ of normalized sum of weighted squared deviations between experimental data not used in the modeling and modeled structures, χ^2^, normalized for a certain confidence level of 68%, vs. the RMSD of the best structure scored by the data not used for modeling. Models below the green horizontal line agree with the validation data. (**c**) Posterior model density, density of validated structures, and reference (ground truth).

## Supplementary Notes

### Supplement Note 1. Forward model for the fluorescence correlation

We use the normalized diffusion term, *G*_*diff*_, for a 3-dimensional (3D) Gaussian shaped detection/illumination volume to describe the fluorescence correlation curves. The term *G*_*diff*_ is given by:

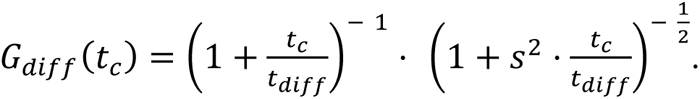

Here, *s* is a shape parameter of the detection volume, *t*_*diff*_ is the diffusion time of the molecule in the detection volume, *t*_*c*_ is the correlation time. The green, G, and red, R, auto and cross correlation function, *G*, were described by:

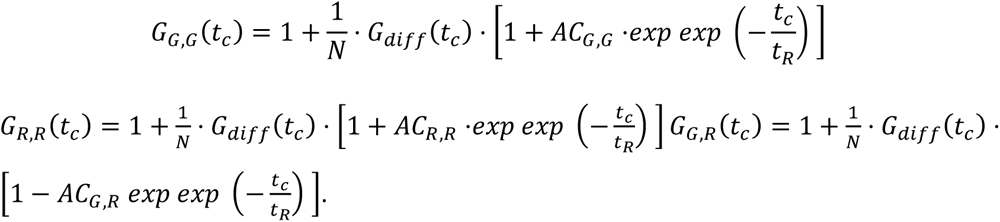

Above, *AC_*G,G*_*, *AC_R,R_*, and *AC_G,R_* are the amplitudes of the kinetic reaction terms and *N* is the effective number of molecules.

### Supplement Note 2. Scoring function in structural modeling

We represent, score, and sample the molecular system using *p*(*R*_*DA*_|*F*, *r*) as surrogate for *F* and *r*. To score structures, we implemented a forward model for IMP that computes: (*i*) the distribution of the fluorophores around their attachment site, (*ii*) inter-dye distance distributions for the fluorophores. We score structures by a sum of weighted squared deviations between the Data, *D*, and the model, *M*:

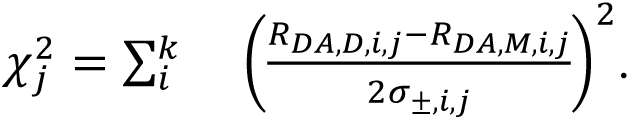

Here, *i* refers to the FRET pair, *k* is the number of FRET pairs, *j* is the state (*C*_1_ or *C*_2_), *R*_*DA, D,i,j*_ and *R*_*DA, M,i,j*_ represent inter-dye distance of the i^th^ FRET pair and the j^th^ state for the data, *D*, or the model, *M*, respectively, and σ_±,*i,j*_ is a weighting parameter for positive and negative deviations for the i^th^ distance of the j^th^ state. Here, *R*_*DA, M,i,j*_ is the average inter-fluorophore distance between for sterically allowed fluorophore positions in a fluorophore pair.

